# Whole-genome sequencing of major malaria vectors revealed an evolution of new insecticide resistance variants in a longitudinal study in Burkina Faso

**DOI:** 10.1101/2023.11.20.567800

**Authors:** Mahamadi Kientega, Chris S Clarkson, Nouhoun Traoré, Tin-Yu J Hui, Samantha O’Loughlin, Abdoul-Azize Millogo, Patric S Epopa, Franck A Yao, Adrien M G Belem, Jon Brenas, Alistair Miles, Austin Burt, Abdoulaye Diabaté

**Affiliations:** Institut de Recherche en Sciences de la Santé (IRSS), Bobo-Dioulasso 01 BP 545, Burkina Faso; Vector Surveillance Programme, Genomic Surveillance Unit, Wellcome Sanger Institute, Hinxton, Cambridge, UK; Université Nazi Boni, Bobo-Dioulasso 01 BP 1091, Burkina Faso; Department of Life Sciences, Imperial College, London SW7 2AZ, UK; Institut des Sciences des Sociétés, Ouagadougou 03 BP 7047, Burkina Faso

## Abstract

Intensive deployment of insecticide based malaria vector control tools results in the rapid evolution of phenotypes resistant to these chemicals. Understanding this process on the genomic level is essential for the deployment of successful interventions. Using whole genome sequencing data of 1409 individual *An. Gambiae* s.l. collected from 2012 to 2017, we investigated the change in genetic structure and the evolution of the insecticide resistance variants in natural populations over time and space. The results showed similar and constant nucleotide diversity and negative Tajima’s D between *An. gambiae* s.s. and *An. coluzzii*. PCA and *F_ST_* showed a clear genetic structure in the *An. gambiae* s.l. species. Genome-wide *F_ST_* and H12 scans identified genomic regions under divergent selection and also having an implication in the adaptations to ecological changes. Novel voltage-gated sodium channel pyrethroid resistance target-site alleles (*V402L, I1527T*) were identified at increasing frequencies alongside the established *kdr* alleles (*Vgsc-L995F*, *Vgsc-L995S* and *N1570Y*) within the *An. gambiae* s.l. populations. Organophosphate metabolic resistance markers were also identified, at increasing frequencies, within the *An. gambiae* s.s. populations from 2012 to 2017, including the SNP *Ace1-G280S* and its associated duplication. Variants simultaneously identified in the same vector populations raise concerns about the long-term efficacy of new-generation bednets and the recently introduced organophosphate pirimiphos-methyl indoor residual spraying. These findings highlighted the benefit of genomic malaria vector surveillance for the detection of new insecticide resistance variants, the monitoring of the existing resistance variants, and also to get insights into the evolutionary processes driving insecticide resistance.

**Author Summary:** Genomic surveillance of malaria vectors is crucial for understanding the genetic variation in natural vector populations and also guiding the implementation of novel and innovative vector control tools. Application of sequencing technologies in vector studies provide insights on the genetic and evolutionary phenomena of vectors that could have impact on vector control strategies. By analyzing the whole genome data of 1409 wild *An. gambiae* s.l. mosquito collected between 2012 and 2017, we showed an emergence of novel insecticide resistance markers alongside increasing frequencies of existing insecticide resistance variants over time in Burkina Faso. We showed the benefit of genomic surveillance of malaria vectors for the monitoring of the insecticide resistance variants and also providing insights into the evolutionary processes driving insecticide resistance.

## Introduction

Malaria remains a major public health problem in Africa [1]. *Anopheles gambiae* s.l., the main malaria vector in Africa, is phenotypically and genotypically diverse, allowing rapid adaptation to environmental changes, such as introductions of new insecticidal control methods [2]. In Burkina Faso, major malaria vectors *An. gambiae* s.s., *An. coluzzii* and *An. arabiensis* can be found living in sympatry, with diverging resting and feeding behaviours [3,4]. These species, through their ability to colonize different ecological settings, their preference for human blood feeding, and their high susceptibility to a malaria parasite (*Plasmodium falciparum*) infection, are responsible for around 90% of the malaria burden in Burkina Faso and the whole sub-Saharan Africa [5,6].

The National Malaria Control Programme (NMCP) of Burkina Faso and their partners have invested much to increase the coverage of long-lasting insecticide treated bednets (LLINs) and other control tools such as indoor residual spraying (IRS) of insecticides to drop the malaria transmission line in the country. The coverage of LLINs in households increased significantly in the country over the years (5.6% in 2003 to about 80% in 2014) reaching 83% in 2021 [7,8]. The recent 2019 and 2022 bednet campaigns involved the free distribution of three types of LLINs, especially the piperonyl butoxide (PBO)-synergist, the dual-AI Interceptor G2 and the standard LLINs [9]. Despite these achievements, malaria remains a countrywide health problem responsible for about 12.2 million cases and 4355 deaths during the year 2021 [10]. In Burkina Faso and most sub-Saharan countries, the primary vector surveillance strategies relied on the routine tracking of the vector species density and distribution and the monitoring of insecticide resistance (IR) status, unable to properly explain the genetic and evolutionary processes driving the increasing spread of IR variants and vector behavioral changes [11]. Recent advances in genomic sequencing technologies, both price and throughput, and new cloud native analysis tools have made genomic surveillance of malaria vectors possible. Genomic surveillance has advantages to provide insights on the genetic and evolutionary phenomena of vectors that could have any impact on vector control strategies. Towards this aim, the *Anopheles gambiae* 1000 genomes (Ag1000G) project driven by the MalariaGEN Vector Observatory sequenced more than 15000 *Anopheles* mosquitoes collected in 25 African countries [12,13]. A major goal of this project was to support malaria elimination efforts by providing a high quality open access data resource on natural genetic variation within *An. gambiae* s.s., *An. coluzzii* and *An. arabiensis* populations, both for the public health and research communities [14].

Previous studies investigating insecticide resistance (IR) using these genomic data have highlighted associated genetic variants distributed throughout the mosquito genome [15–18]. The phenotypic effect of many of these variants has been demonstrated through in-vivo and in-vitro experiments and their effects are evident in low mortality rate observed during the IR bioassays, and in the reduction of the efficacy of the insecticide-based control tools [19–21]. The strength of these resistance phenotypes and the wide geographical distribution of these variants raise concerns about the durability of the new insecticide-based vector control tools being used to overcome resistance, including indoor residual spraying (IRS) with organophosphate insecticides such as Actellic (pirimiphos-methyl) and pyrethroid bed-nets treated with metabolic resistance blocking piperonyl butoxide (PBO) synergists.

There is an increasingly urgent need for sustainable strategies and effective tools to support malaria elimination efforts in the face of an ever evolving vector. Research of alternative and innovative tools such as genetic engineering, new insecticidal compounds as well as natural symbionts has increased in recent years, giving a glimmer of hope to people living in endemic areas [22,23]. However, given the genome complexity of the malaria vectors, highlighted by many studies [12,13,24], any implementation of vector control tools requires an understanding of the genomic variation of vector species, particularly an understanding of the evolutionary processes, emergence and the spread of the variants associated with IR in natural populations over time. Thus, longitudinal sampling of vectors, followed by whole genome population genomics becomes an essential tool upstream and downstream of the implementation of the control tools to provide insights on the genetic and evolutionary phenomena of vectors that could have any impact on vector control strategies. Here we leverage the 1409 *Anopheles gambiae* s.l. samples, collected in Burkina Faso between 2012-2017 and sequenced as part of the *Anopheles gambiae* 1000 Genomes Project (2017, 2020), to characterize the spread of known IR variants over space and time, while analyzing these natural populations for evidence of the evolution of novel IR mechanisms which could threaten current or future vector control methods.

## Results

### Population Sampling

Following the *Anopheles gambiae* 1000 Genomes project, further mosquito collections were sequenced as part of the follow-up project, the MalariaGEN Vector Observatory. In total, this resulted in 1409, deep, whole-genome sequenced, mosquitoes collected from three villages surrounding Bobo-Dioulasso city in western Burkina Faso. Mosquitoes were collected longitudinally, across the years 2012, 2014, 2015, 2016 and 2017 (Fig 1), though species representation varies between collection sites and time-points (Table 1). *An. coluz*zii was the most predominant species found, accounting for 52.73% [743/1409] of samples, followed by *An. gambiae s.s*. with 38.96% (549/1409), while *An. arabiensis* only comprised 8.16% (115/1409) of samples collected. Two *An. gambiae* s.s. and *An. coluzzii* hybrids were also identified. As shown by previous work, *An. coluzzii* remains the most predominant species in Bana (83.87% [489/583]) and Souroukoudinga (60.90% [243/399]) while *An. gambiae* s.s. is prevalent in Pala (70.72% [302/427]) [6].

**Fig 1.**
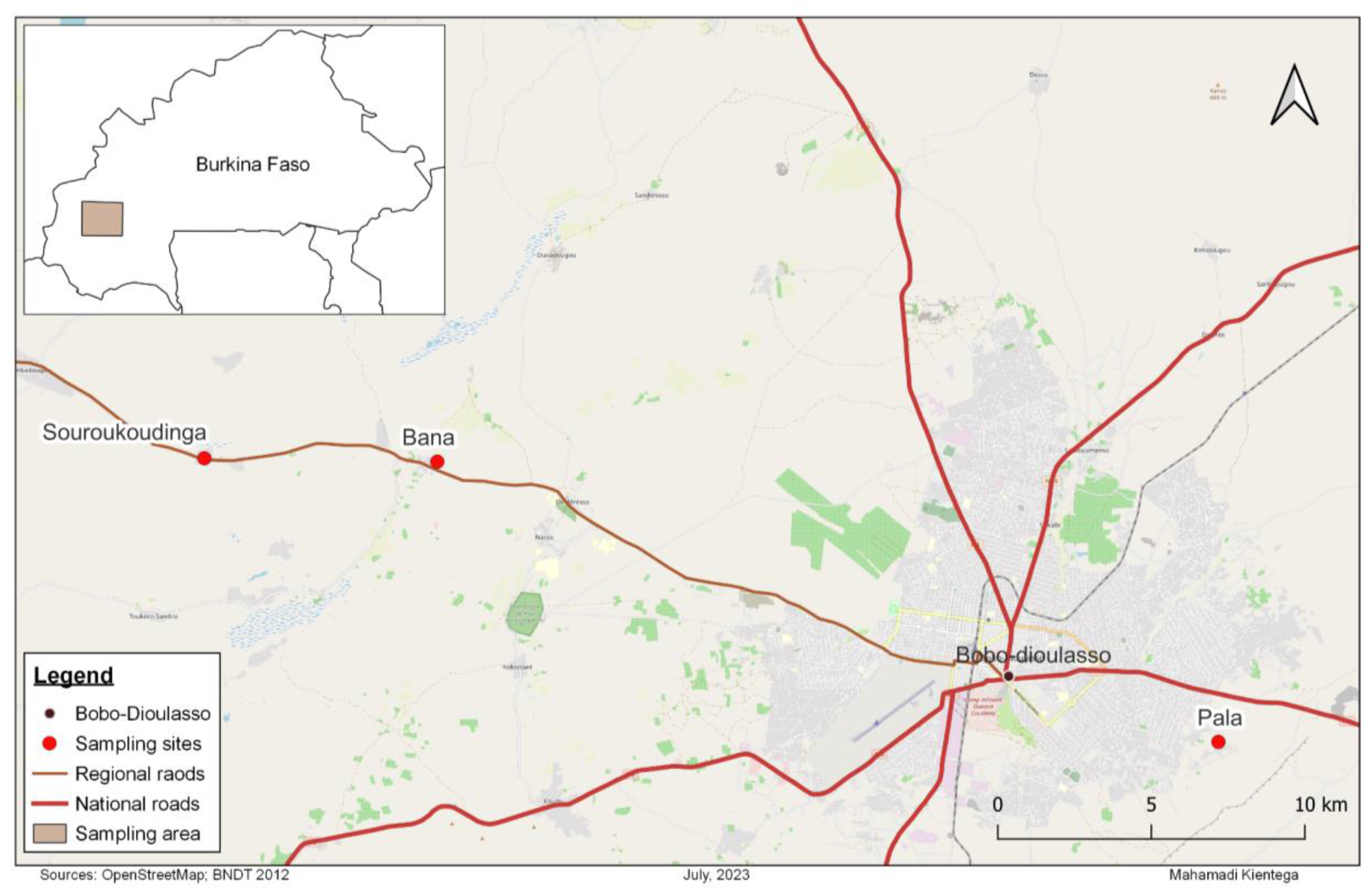
Map of the sampling sites around Bobo-Dioulasso city

**Table 1.** Species composition of *An. gambiae* complex in the three villages surrounding Bobo-Dioulasso city over the five collection years.

| Species | Periods | Sampling sites |  |  |
| --- | --- | --- | --- | --- |
|  |  | Bana | Souroukoudinga | Pala |
| *An. gambiae* s.s. | 2012 | 22 [33.85%] | 28 [49.12%] | 48 [81.36%] |
|  | 2014 | 32 [24.81%] | 49 [49.49%] | 89 [89.%] |
|  | 2015 | 25 [20.66%] | 40 [31.5%] | 76 [46.34%] |
|  | 2016 | 13 [9.77%] | 38 [44.19%] | 53 [79.1%] |
|  | 2017 | 0 | 0 | 36 [97.3%] |
| *An. coluzzii* | 2012 | 42 [64.62%] | 29 [50.88%] | 11 [18.64%] |
|  | 2014 | 96 [74.42%] | 50 [50.51%] | 0 |
|  | 2015 | 96 [79.34%] | 87 [68.5%] | 0 |
|  | 2016 | 120 [90.23%] | 48 [55.81%] | 0 |
|  | 2017 | 135 [100.%] | 29 [96.67%] | 0 |
| *An. arabiensis* | 2014 | 1 [0.78%] | 0 | 11 [11.%] |
|  | 2015 | 0 | 0 | 88 [53.66%] |
|  | 2016 | 0 | 0 | 14 [20.9%] |
|  | 2017 | 0 | 1 [3.33%] | 0 |
| *intermediate_gambiae_c*<br>*oluzzii* | 2012 | 1 [1.54%] | 0 | 0 |
|  | 2017 | 0 | 0 | 1 [2.7%] |

### Genetic Diversity

Understanding genetic diversity allows us to compare demography of species over time and space. Analyses of these data showed around 27.94 million of segregating sites (∼ 91% of biallelic SNPs) in the *An. gambiae* s.l. genome. Approximately 25.68 million of SNPs were identified in the autosomes and 2.262 million of SNPs in the X chromosome. Analysis to evaluate the genetic diversity of this variation (see Methods), demonstrated that nucleotide diversity (*θ_π_*) and Watterson’s theta (*θ_w_*_)_, were high and stable over all collection years for *An. gambiae* and *An. coluzzii*, similar to the majority of natural populations across the species range (Ag1000g, 2017) (Fig 2), potentially suggesting similar demographic histories between these two species. In contrast, *An. arabiensis* showed lower diversity, though with apparent increase over collection years. The diversity findings were reflected in a genome-wide negative Tajima’s D, caused by an excess of rare variants due to either positive selection and/or rapid demographic changes through population expansion. In addition, the allelic frequency spectrum (Fig S1 & S2) also showed the same trend highlighting the excess of low frequency variants in the *An. gambiae* s.l. genome.

**Fig 2.**
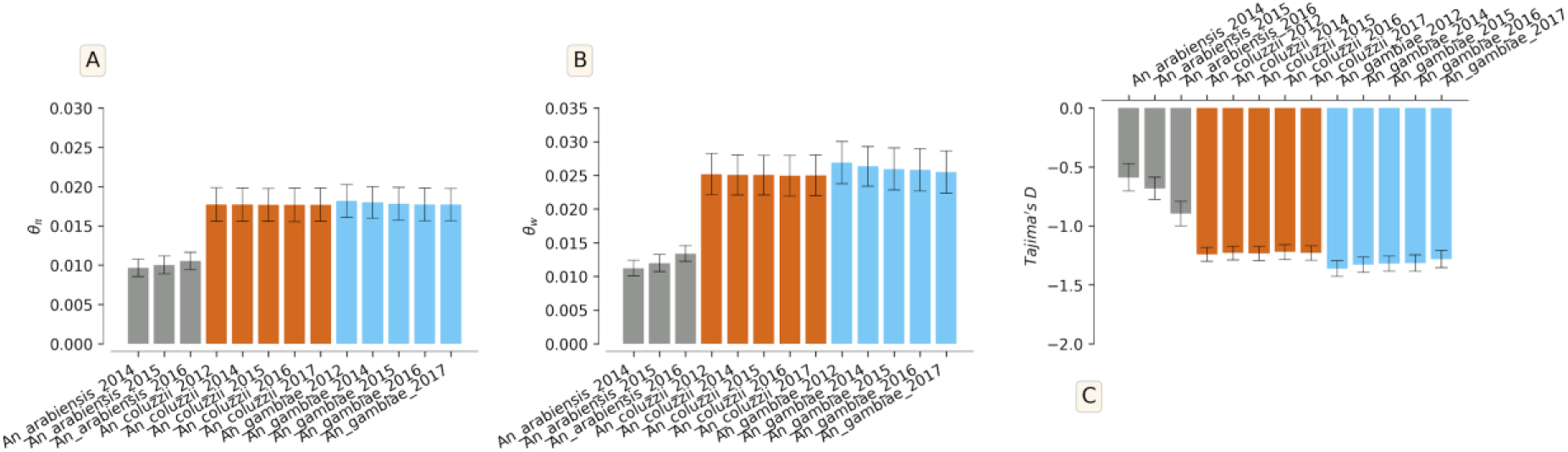
Average of the diversity statistic (A: nucleotide diversity (*θ_π_*), B: Watterson theta (*θ_w_*), C: Tajima’s D) in the *An. gambiae* s.l. populations

### Population Structure

Understanding population structure between, and within, vector species is not only important to understand due to its implications in gene flow of medically relevant alleles or role out of gene drive control methods, but hidden structure will confound downstream population genomic analyses. Consequently, the genetic structure and the patterns of gene flow were investigated using SNPs on the 3L chromosome (see methods). Pairwise *F_ST_* was estimated between species collected in different villages and was found to be low between *An. gambiae* and *An. coluzzii* scaling from 0 to 0.05 (Fig 3A). These results suggest gene flow between *An. coluzzii* and *An. gambiae* s.s.. *An. arabiensis*, however, showed relatively high *F_ST_* between both *An. coluzzii* and *An. gambiae*, ranging from 0.26 to 0.30 (Fig 3A), potentially suggesting weaker gene-flow between species. No differentiation (*F_ST_* ∼ 0) was observed between populations of the same species collected in different villages, suggesting no population structure on this scale.

**Fig 3.**
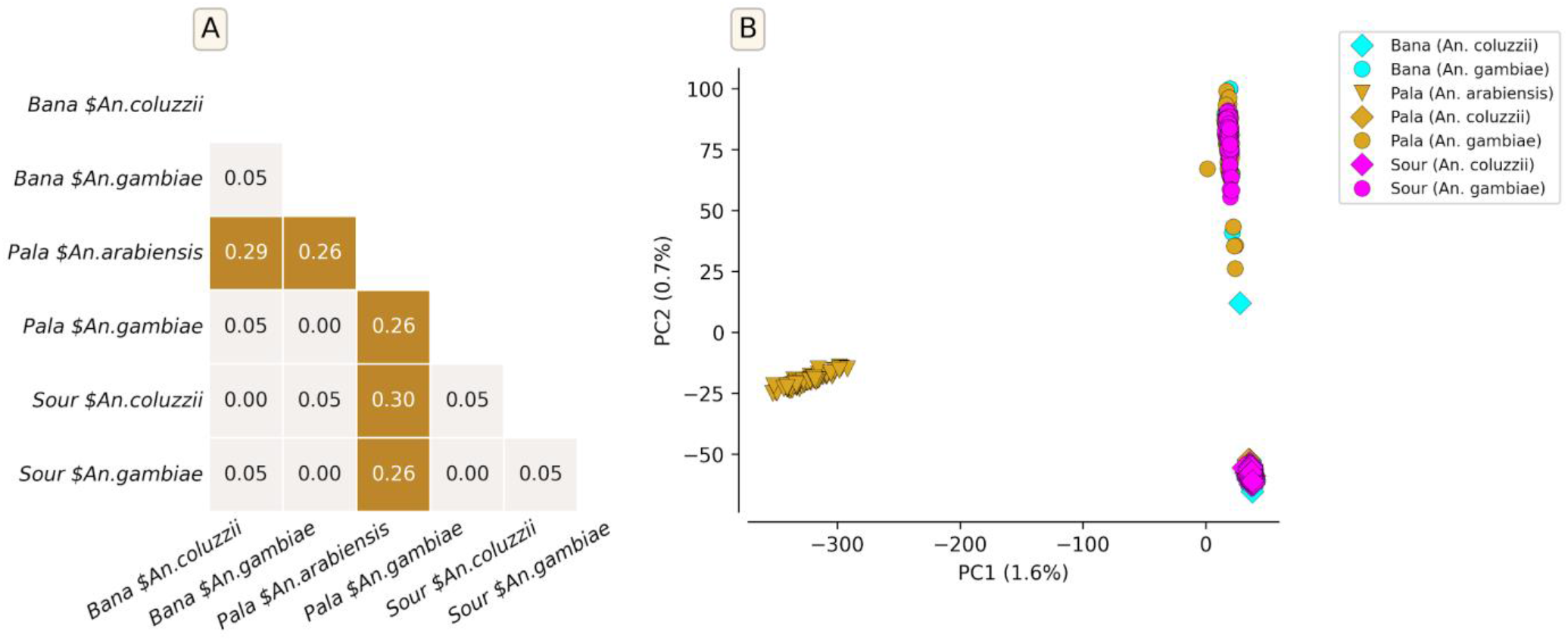
Genetic differentiation and population structure of *An. gambiae* s.l. using the SNPs identified in the 3L chromosome. A: Pairwise *F_ST_* between *An. gambiae* s.l. populations collected in different villages. No differentiation between populations of the same species collected in two different villages. B: PCA showing the genetic structure of *An. gambiae* s.l. populations.

Principal component analysis (PCA) showed a clear divergence between the *An. gambiae* s.l. species. Examining the first two principal components, samples fell into three clusters, where individuals of the same species were more genetically related than to others. No sub-clusters were observed within each species (Fig 3B). No change was observed in the population structure of each species throughout the years, and the sampling sites (Fig S3); however, some other genomic regions were shown to be under divergent selection (see Organophosphate and carbamate resistance: increasing threats to control plus a potential novel IR mechanism, Fig S4). The lack of genetic structure (low *F_ST_ ∼ 0.05*) between *An. coluzzii* and *An. Gambiae* s.s. showed that these species are capable of sharing advantageous variants in this region Burkina Faso, such those involved in insecticide resistance.

### Insecticide Resistance

#### Pyrethroid Target-site Resistance: concerning increase of new mutations

We investigated the distribution of genetic variants in the voltage gated sodium channel gene (*Vgsc)*, which codes for the target-site protein of the pyrethroid insecticides used in bednets). Non-synonymous SNPs in insecticide target site genes, can lead to insecticide resistance via physically altering the conformation of the target-site protein, causing reduced insecticide binding [25].

Pyrethroid target-site resistance in *Anopheles gambiae* s.l. has, historically, been primarily driven by two main “*kdr”* SNPs, causing the amino acid changes *Vgsc-L995F* and *Vgsc-L995S* [17]. These mutations are widespread, and are at high frequencies in many regions in Africa [12]. Improvements in sequencing technologies have allowed the identification of many additional SNPs in the *Vgsc* gene that could enhance the level of pyrethroid resistance over the time [26,27]. This study identified 634 non-synonymous SNPs in all the *An. gambiae* s.l. populations (Table S1). The *kdr* allele *Vgsc-L995F* was identified at high frequencies (> 30%) in all the populations, and was found at fixation in *An. gambiae* s.s. collections from 2012. While *An*. *arabiensis* populations also showed high frequencies of *Vgsc-L995S* (Fig 4, Fig S5). The *Vgsc-N1570Y* allele, previously shown to increase the level of pyrethroid resistance when in concert with *Vgsc-L995F* [26], was also present in both *An. gambiae s.s*. and *An. coluzzii* populations at around 30% frequency. Some SNPs were found to be species specific over the years. We identified the alleles *Vgsc-V402L, N1345I, I1527T, K1603T, L1667M, P1874S, A1934V* and *I1940T* specifically in *An. coluzzii* populations whereas *Vgsc-V1853I, I1868T, E1597G* were specific to *An. gambiae* s.s. populations (Fig S5). Most of these species specific alleles were identified at low frequencies.

**Fig 4.**
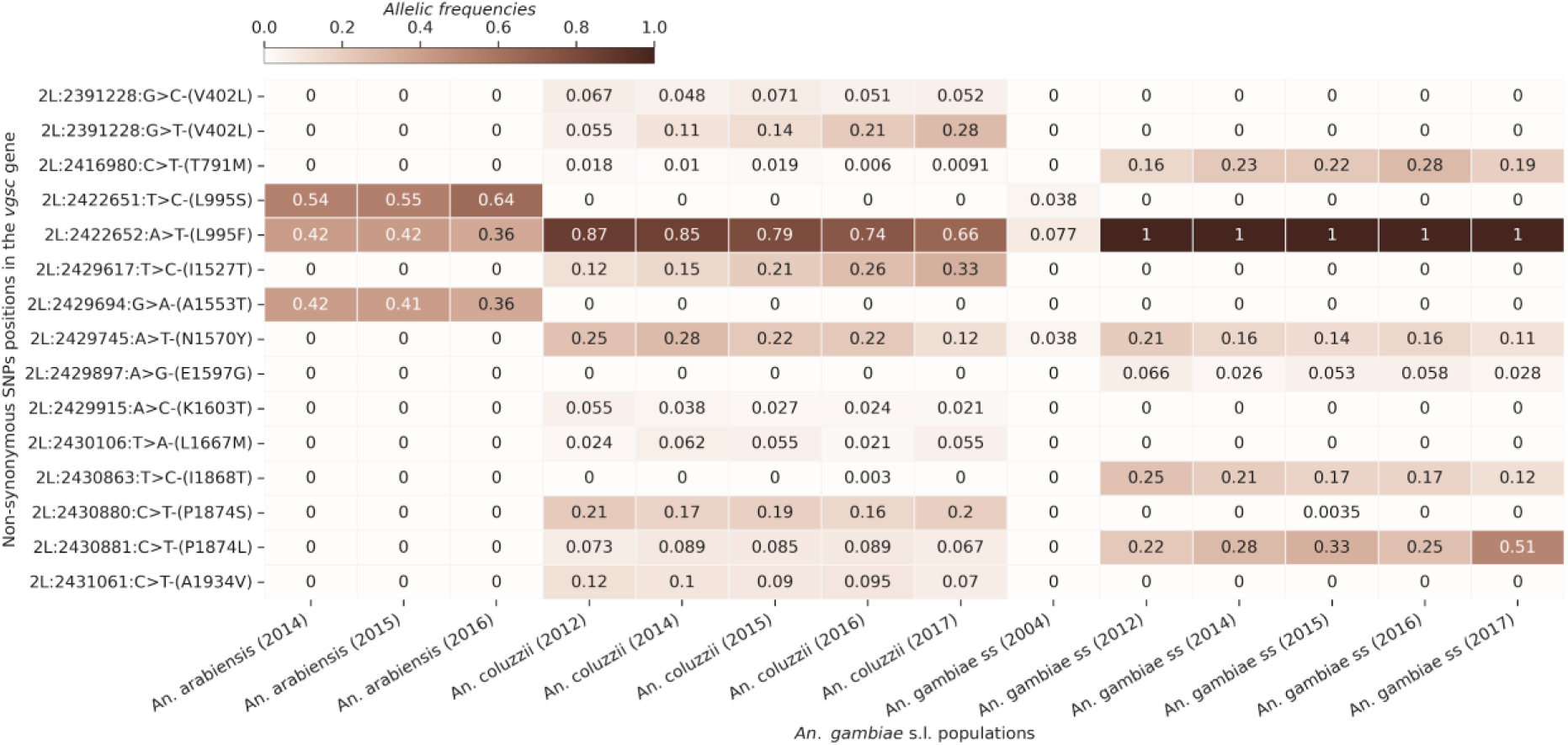
Heat map showing the frequencies of the non-synonymous SNPs frequencies (max freq > 0.05) in the *Vgsc* gene in the *An. gambiae s.l.* populations over time. The X axis shows the *An. gambiae s.l.* populations and the years of collection. The Y axis shows the non-synonymous SNPs positions in the chromosome 2L. The gradient color bar shows the distribution of the allelic frequencies.

A *Vgsc* haplotype carrying the apparently linked amino acid substitutions *Vgsc-V402L+I1527T* was also identified in Burkina Faso *An. coluzzii* (Fig 4 and 5, Appendix 5). A recent *in-vivo* investigation has shown that the *Vgsc-V402L* substitution results in a pyrethroid resistance phenotype with lower fitness costs than the classic *Vgsc-L995F kdr* mutation [27]. Upon discovery of *Vgsc-V402L+I1527T,* it was suggested that this haplotype may be a “relic” resistance haplotype, pre-dating and being overtaken by a more effective *Vgsc-L995F* carrying haplotype [17]. However, our time series data shows that the inverse appears to be the case. *Vgsc-V402L+I1527T* increased from 12.19% in 2012 to 33.12% in 2017 (Fig 4 and 5, Fig S5). The increase of these alleles coincides with the drop in the frequency of the *Vgsc-L995F* mutation in *An. coluzzii* populations from 86.58% in 2012 to 66.46% in 2017 (Fig 4 and 5). Our results and other works [17,27] suggest these two alleles are increasing in frequency and replacing *Vgsc-L995F* in *An. coluzzii*, but in 2017 had not spread to *An. Gambiae* s.s. The increased emergence of new *kdr* alleles alongside the existing *kdr* mutations (*Vgsc-L995F* and *Vgsc-L995S*) at high frequencies may increase the resistance level of vector populations to pyrethroid compounds.

**Fig 5.**
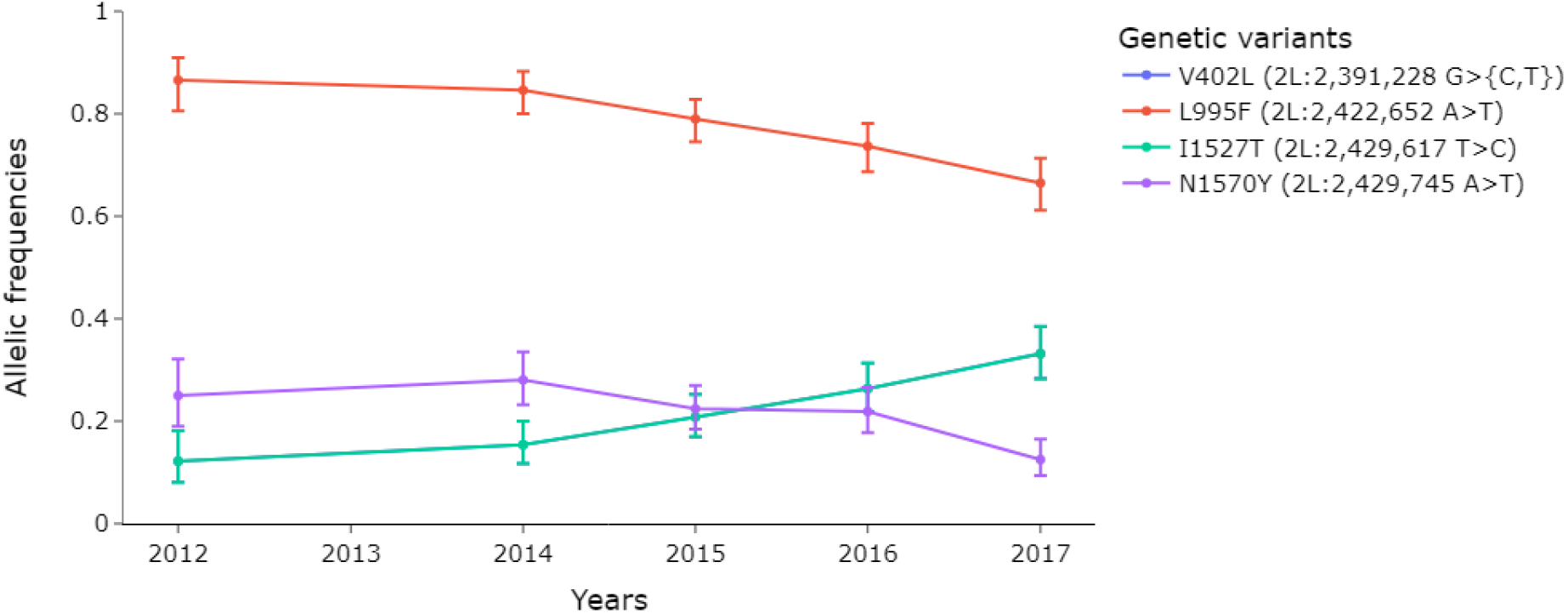
Dynamics of *Vgsc-L995F* and double mutant *Vgsc-V402L+I1527T* allelic frequencies in the *An. coluzzii* populations.

#### Pyrethroid Metabolic Resistance: increasing CNV frequencies

Gene duplications and deletions (copy number variants - CNVs) are known to influence the expression level of genes by providing additional copies of the original gene. Consequently, CNVs of genes involved in xenobiotic detoxification can increase insecticide metabolism, producing a resistance phenotype [15]. To help overcome pyrethroid resistance, now commonplace in many natural populations, PBO synergist bednets have recently been introduced. These bednets enhance the pyrethroid efficacy by inhibiting the cytochrome P450 enzymes that can be responsible for metabolic pyrethroid resistance [28]. Often, mechanisms driving the resistance in a given area remain unclear [20], however, whole genome sequencing allows CNV calling, and the detection of resistance associated alleles [16].

CNVs were identified in 136 P450 genes distributed across the whole genome and most of these (∼ 63.97 %) were gene amplifications (Table S2). The majority of the CNVs were species-specific, and some were found at increasing frequencies over time in the three species. Some genes showing low CNVs frequencies in 2012 displayed increasing CNVs frequencies over the time (Fig 6). This situation was observed in the *Cyp6z* cluster of genes (*Cyp6z1, Cyp6z2 and Cyp6z3*), known to be associated with pyrethroid resistance [29], whose CNVs frequencies increased from 17% in 2012 to up to 60% in 2017 in *An. gambiae* s.s. populations. Additionally, the *Cyp9k1* gene, also previously shown to be involved in pyrethroid resistance, showed CNVs at high frequencies up to 80% from 2012 to 2017. In the *An. coluzzii* populations, CNVs were identified at frequencies ranging from 50% in 2012 to up to 90% in 2017 within the *Cyp6aa1, Cyp6aa2, Cyp6p15p* and *Cyp12f2* genes. Moreover, the *Cyp6af* (*Cyp6af1* and *Cyp6af2*) and *Cyp9m* cluster genes (*Cyp9m1 and Cyp9m2*) exhibited high CNVs frequencies (50 – 100%) in all the populations (Fig 6).

**Fig 6.**
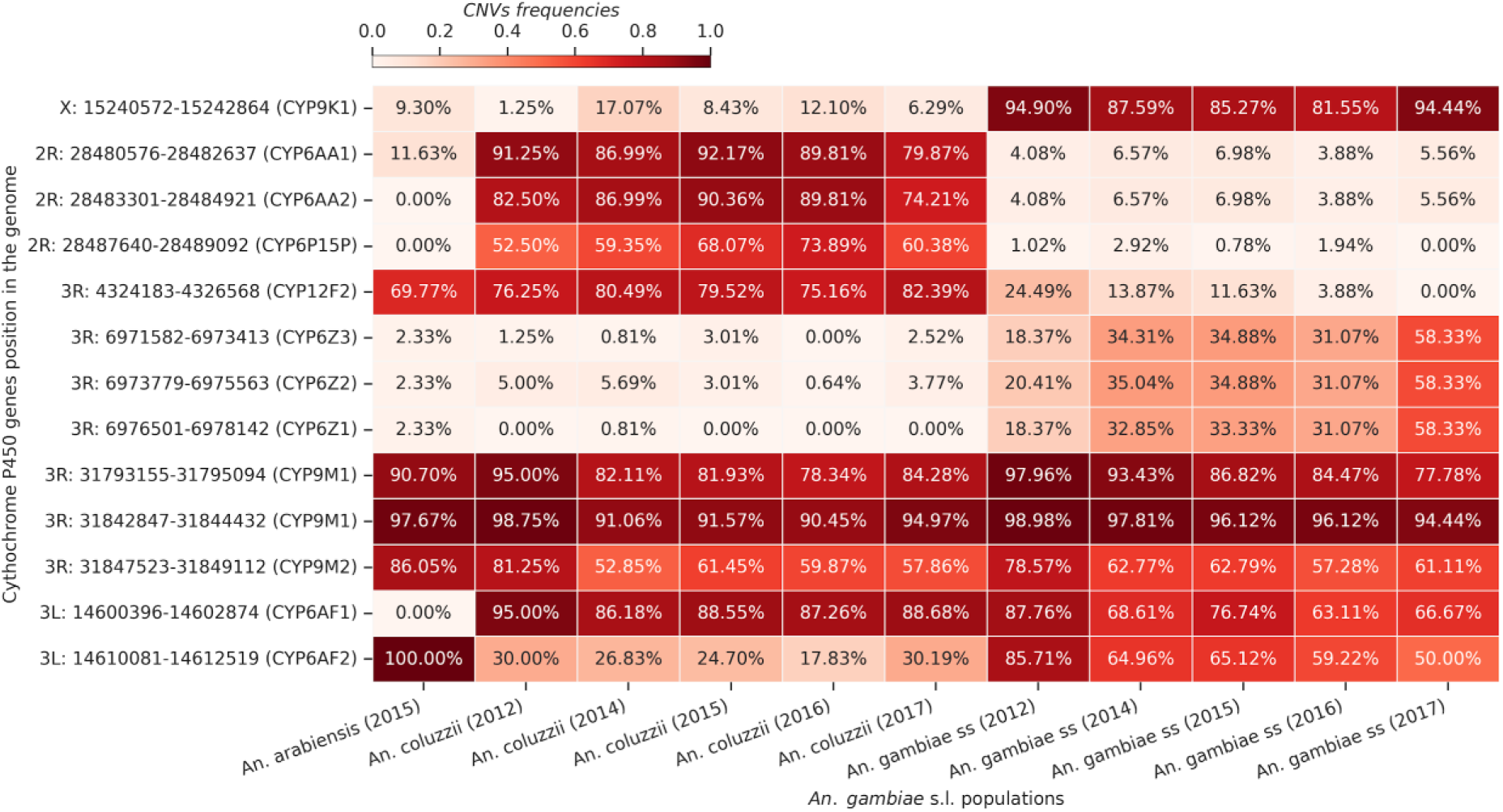
Heat map of the cytochrome P450 genes showing high CNVs frequencies in the *An. gambiae* s.l. populations. The X axis shows the *An. gambiae* s.l. populations and the years of collection. The Y axis shows the positions of the P450 genes in the genome. The gradient color bar shows the distribution of the allelic frequencies.

#### Organophosphate and carbamate resistance: increasing threats to control plus a potential novel IR mechanism

Indoor residual spraying (IRS) remains another major vector control strategy in Africa [30]. However, the coverage of this strategy dropped considerably over the years because of the emergence of resistance to the pyrethroid compounds and also the low residual efficacy of carbamates compounds [31]. Recently, IRS with Actellic 300 CS containing pirimiphos-methyl, an organophosphate as the active ingredient, showed interesting results including a prolonged residual efficacy up to seven months and a reduction of vector density in many sprayed areas in Africa [32–34]. This new formulation demonstrated better ability to target pyrethroid resistant and indoor resting mosquitoes and showed additional benefit when used in combination with LLINs [35,36].

Using the time series collection, we investigated the magnitude of resistance to organophosphates and carbamates resistance in Burkina Faso that could threaten the long term efficacy of pirimiphos-methyl-based IRS. A sliding window genome-wide *F_ST_* scan showed a year to year (from 2012 to 2017) genetic difference in some regions of the genome especially on chromosomes 2 and X in *An. gambiae* s.s. and *An. coluzzii* samples. Most of these changes were observed in genomic regions previously shown to be associated with insecticide-resistance suggesting a signal of positive selection in these genes (Fig 8, 9 and 10).

Both *Ace1* and Diacylglycerol Kinase (DGK) loci showed no selection signal in 2012 in any vector populations we sampled, but by 2017 selection signals were detected at both loci in *An. gambiae* s.s. populations. The increasing gene amplification signal in the *Ace1* locus, from 32.65% in 2012 to up to 91.67% in 2017, was corroborated by the increase in the insecticide resistance associated *Ace1-G280S* variant previously shown to be involved in organophosphate and carbamate resistance [37] (from 15.81% in 2012 to 65.27% in 2017), but only in *An. gambiae* s.s. populations (Fig 7). We also identified 243 other non-synonymous SNPs in the *Ace1* gene (Table S3). CNV amplifications were also found in *An. coluzzii* populations, but at low and constant frequencies (∼ 3%), but not in *An. arabiensis* populations.

**Fig 7.**
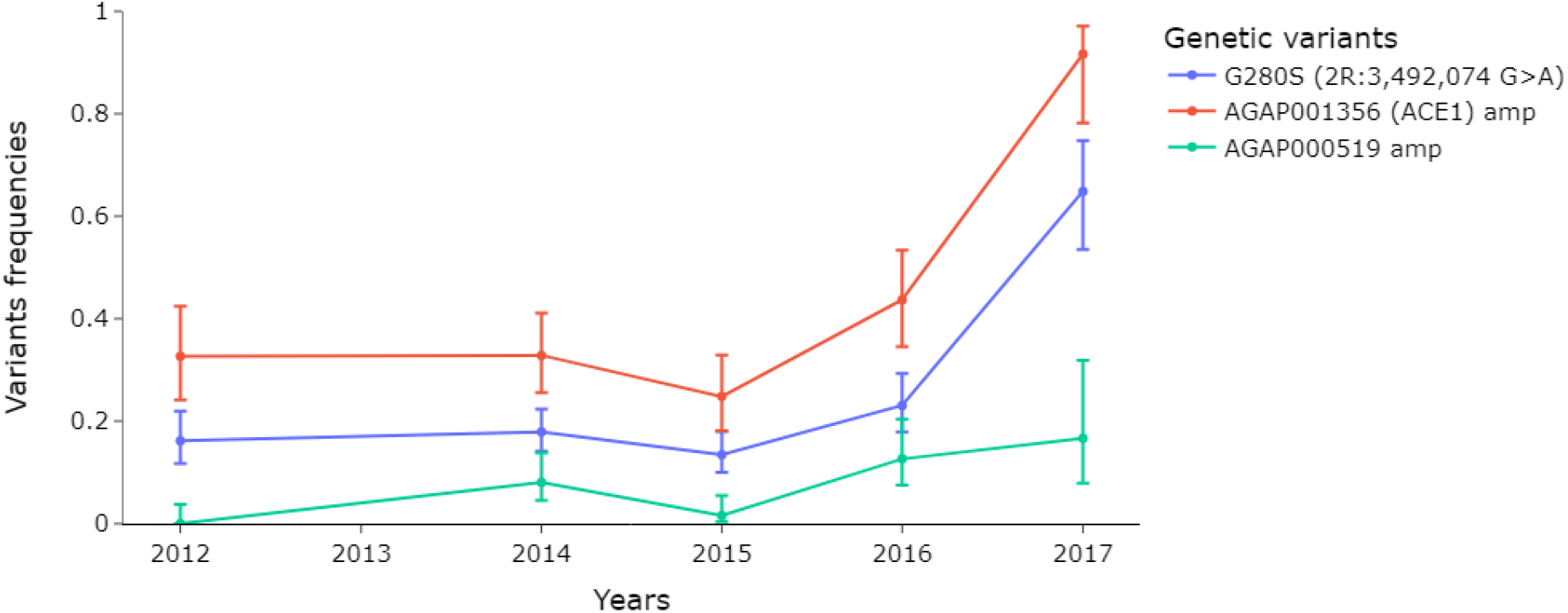
Evolution of the *Ace1* and Diacylglycerol Kinase (AGAP000519) CNV frequencies in the *An. gambiae* s.s. populations.

Genetic variants of the *Gste* genes have also been reported to be involved in resistance to organophosphates [38]. In this study, we identified CNVs in 41 genes (24 genes with amplifications and 17 with deletions) of the *Gste* genes. These CNVs are distributed across the genome (Table S4). Most of these CNVs were found at low frequencies (< 50%) except the *GstD5* which showed high CNV frequencies (60% to 100%) in all the populations (Fig S6, Table S4).

The carboxylesterases (COEs) enzymes have been reported to be involved in the rapid metabolization of both pyrethroid and organophosphate compounds [39] and consequently could have an impact on the benefit of the combined organophosphate-based IRS and pyrethroid-based LLINs approach. In this study, CNVs were identified in 39 genes across the genome, most of these CNVs were gene amplifications, only 25.64% were deletions (Table S5). The frequencies of these CNVs were relatively low, except those of the *Coeae*60 gene which were identified at frequencies above 60% in *An. coluzzii* populations from 2012 to 2017 (Fig S7). Additionally, a positive selection signal was also detected in the *Coeae*60 gene (Fig 8). The *Coeae*G* cluster genes (*Coeae2G, Coeae3G, Coeae5G, Coeae6G* and *Coeae7G*) and the *Coeae3H* gene showed CNVs at increasing frequencies from 11% in 2012 to up to 25% in 2017 in the *An. gambiae* s.s. populations. The *Coejhe4E* gene showed CNVs at 32.56% frequencies in *An. arabiensis*, and also at frequencies higher than 30% in *An. Gambiae* s.s. populations (Fig S7).

**Fig 8.**
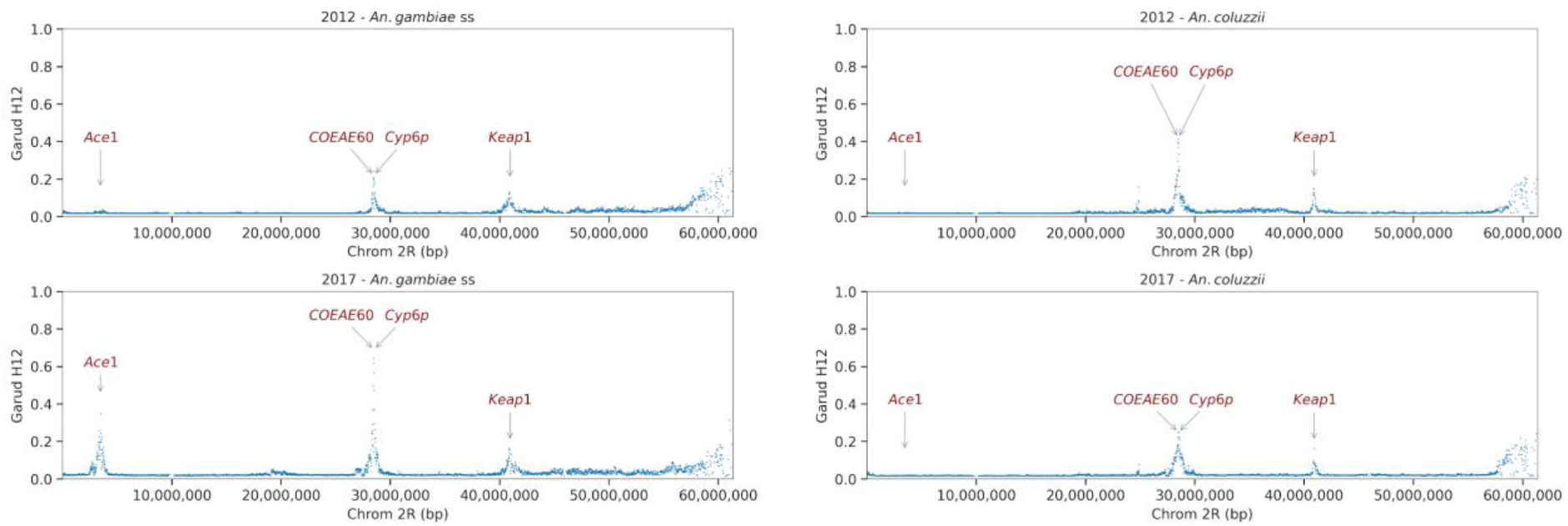
Evolution of the Garud H12 in the chromosome arm 2R of the *An. gambiae s.l.* populations from 2012 to 2017. High values of H12 indicate a signal of positive selection in the corresponding genomic region within the populations: *Ace1: Acetylcholinesterase gene, Cyp6p: Cytochrome P450 gene, COEAE60: Carboxyl-esterase gene, Keap1: Kelch-like ECH-Associated Protein 1*.

The emergence of new forms of insecticide resistance may impact vector control strategy in unexpected ways. Recent studies reported new forms of resistance, especially behavioural changes (early and/or outdoor biting) in *An. gambiae* mosquitoes, in response to the intense use of LLINs causing residual malaria transmission [40,41]. Here, we detected a signal of positive selection in the DGK locus and also an increased frequency of CNVs of this gene within the *An. gambiae s.s.* populations (Fig 9 and 10). No selection signal was detected in vector populations in 2012, with no CNVs identified either. However, in 2017, a signal of positive selection was detected in the DGK locus, with an associated increasing frequency (from 0.00 in 2012 to 16.67% in 2017) of CNVs in vector populations (Fig 7, 9 and 10). Considering this evolution pattern of the DGK gene and also its clear role in the adaptation of *C. elegans* and *Drosophila sp* to environmental change [42,43], this gene may play an important role in the modulation of *An. gambiae* biology in response to the presence of vector control tools.

**Fig 9.**
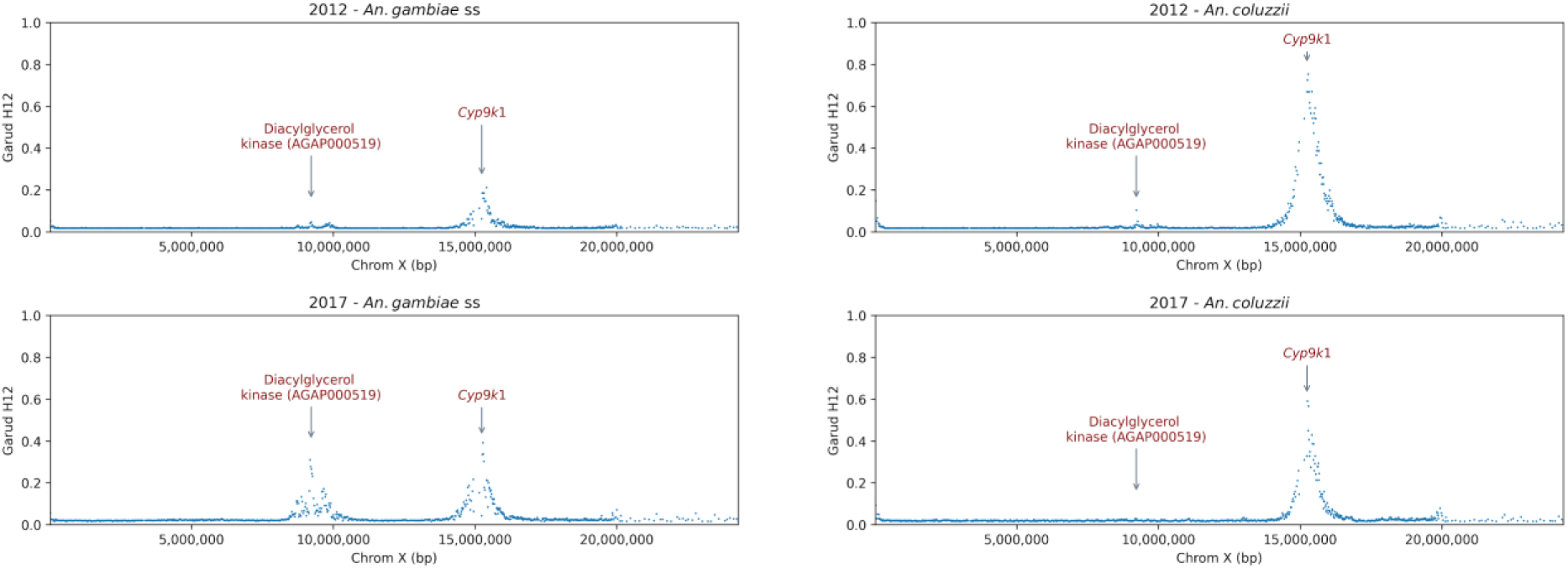
Garud H12 statistic showing a signal of positive selection in the *DGK* and the *Cyp9k1* genes within the *An. gambiae* s.l. populations.

**Fig 10.**
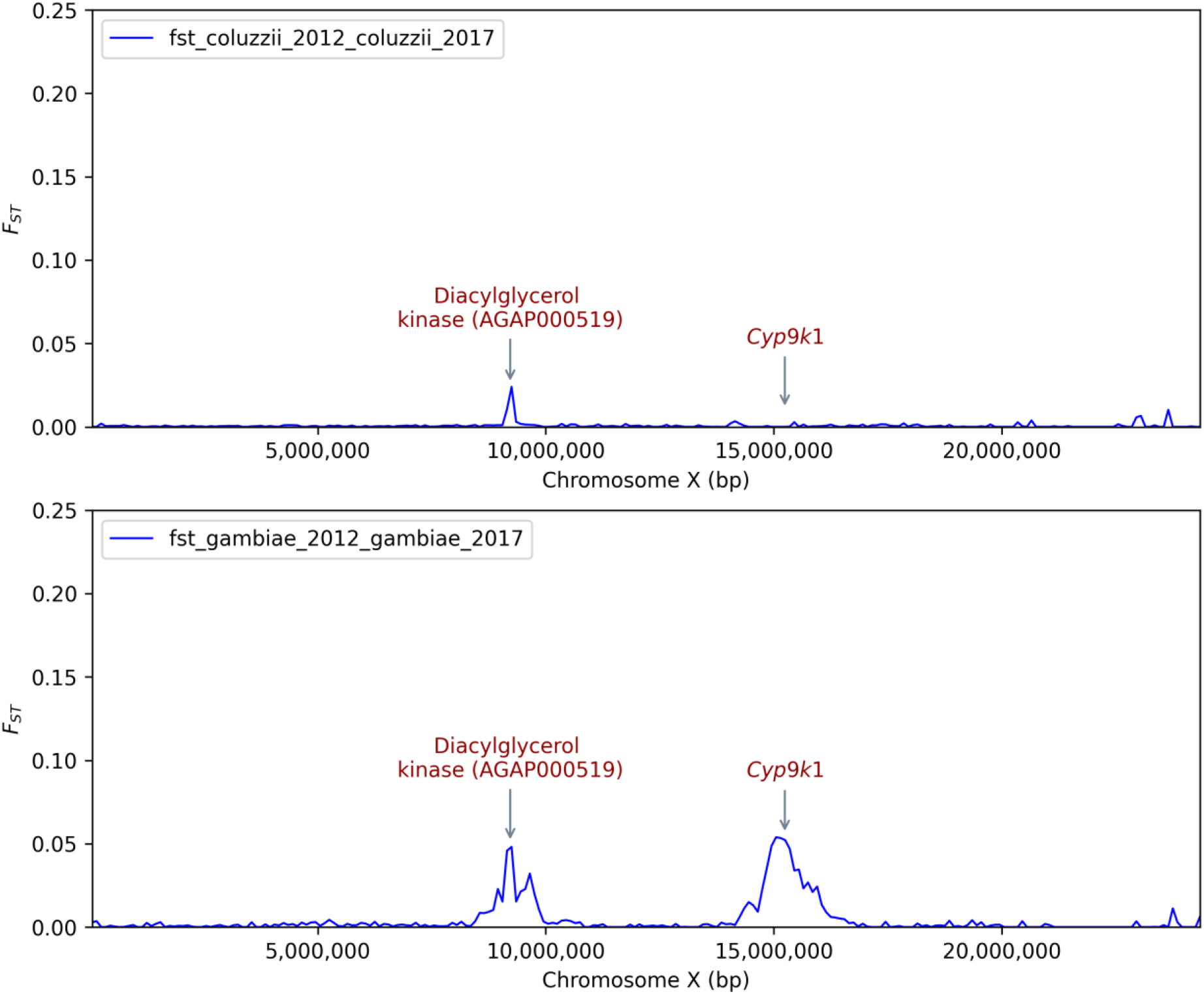
Year to year variation of the genetic differentiation between populations of the same species of *An. gambiae* complex collected in 2012 and in 2017. Signals of differentiation were observed in some regions involved in insecticides resistance including *Cyp9k1* and DGK gene.

## Discussion

Significant advances have been made in the reduction of malaria burden using LLINs and IRS, but the efficacy of these tools is at risk due to the high adaptability of the vector populations to the anthropogenic ecological changes [30]. A better understanding of the molecular, ecological and evolutionary processes allowing vector populations to adapt to the environmental changes would help in the development of innovative tools to support the current control tools in malaria elimination. In this study, we used whole genome sequencing data to study the genetic diversity, the population structure and the evolution of insecticide resistance variants in the *An. gambiae* s.l. populations in the western part of Burkina Faso.

### Bednets – evidence and impacts of target-site and metabolic pyrethroid resistance

The insecticides pressure on malaria vectors lead to an emergence of new insecticide resistance variants that confer fitness benefits to vector populations [44]. This seems to be consistent and evident for the SNPs *Vgsc-V402L+I1527T* which were identified at increasing frequencies in *An. coluzzii* populations. The rapid rise of their frequencies in Burkina Faso *An. coluzzii* populations may be expected given the results of a recent study which revealed the *Vgsc-V402L* produces a pyrethroid resistance phenotype with reduced fitness costs, when compared with *Vgsc-L995F* [27]. The simultaneous emergence of different types of genetic variants around genes and gene clusters known to be associated with pyrethroid metabolism and target site mutations within the *An. gambiae* s.l. populations raise major concerns in the long term efficacy of new pyrethroid insecticide-based control strategies such as LLINs.

Insecticides resistance has been widely studied in Africa, and recent works have reported multiple resistance variants spreading in vectors populations, jeopardizing NMCP efforts in control and elimination [17,18,45]. The implementation of LLINs next-generation bed nets incorporating piperonyl butoxide (PBO) aims to strengthen the efficacy of LLINs by inhibiting the cytochromes P450 enzymes responsible for the rapid detoxifying of pyrethroid insecticide [46]. The high expression of the P450 enzymes was shown to increase the pyrethroid resistance and pre-exposure to PBO restores the susceptibility of malaria vector to pyrethroid [28]. In most cases, the main genetic variants responsible for the insecticide resistance in a given area remain unclear because of the high heterogeneity of the genetic markers mediating metabolic resistance. Although PBO inhibits the cytochrome genes, the emergence of additional new variants could enhance the level of pyrethroid resistance impacting the efficacy of the new vector control tools.

### IRS - evidence and impacts of organophosphates resistance

Another major concern related to this multi-modal resistance is its threat on the effectiveness of the indoor residual spraying (IRS). This strategy is efficient when it is used in combination with LLINs because of its ability to target indoor resting pyrethroid resistant vectors [47]. Although the SNP *Ace1-G280S* has long been identified in *An. gambiae* populations, its frequencies remained low in the populations probably due to its fitness cost in association with the kdr mutation [48]. Indeed, the duplication of the *Ace1* gene (*ace1^D^*) seems to have completely solved the problem of genetic cost yielded by the SNP *Ace1-G280S* [49]. Recent reports showed a wide distribution of this duplication in West African *An. coluzzii* and *An. gambiae* s.s. populations [18,49] probably in response to the use of organophosphate and carbamates compounds in vector control. Similarly, the duplication of *Ace1* gene follows the same trend as the SNP *Ace1-G280S* raising up in frequencies in *An. gambiae* s.s. populations in our study. This clearly showed the close related link between these two variants in conferring high fitness to the mosquito populations. In addition, the CNVs identified at high frequencies in some of the *Gste* and COE genes could increase the expression level of these genes. The simultaneous presence of these variants (CNVs and SNPs) in *Ace1*, *Gste* and esterases genes within the same vector populations strongly contribute to increase the survival capacity of the vector against organophosphates and carbamates. This situation threatens the long-term efficacy of vector control interventions including the recently introduced pirimiphos-methyl-based IRS.

### Whole genome surveillance enables detection of potential novel IR loci

Genome-wide scans for positive selection have proved their usefulness in the identification of genomic regions under divergent selection. In malaria vectors, under high insecticide mediated evolutionary pressure, these regions often have implications in insecticide resistance [50]. We detected a signal of positive selection in a genomic region of the X chromosome corresponding to the DGK gene. Though there is currently no evidence of this gene’s association with insecticide resistant phenotypes in malaria vectors, studies in *Drosophila melanogaster* and *Caenorhabditis elegans* showed its potential involvement in the adaptation to environmental changings [42,43]. In *D. melanogaster,* the rdgA gene, an orthologue of the DGK gene, is eye-specific, strongly involved in light signaling. Any genetic variant influencing the functioning of this gene would probably affect its sensitivity to light, as reported previously [42,51]. Consequently, if light signaling is regulated the same way in mosquito species, the DGK gene may also be involved in the regulation of *An. gambiae* vision. In *An. gambiae*, a previous study showed a rhythmic expression of the genes involved in light signaling under the control of the circadian clock [52]. The expression pattern of these genes suggests their role in calibration of the mosquito visual system to start the nocturnal activities such as flight and biting. In response to the high coverage of LLINs, major changes were reported in vector biting behavior particularly early biting and later biting activities in many areas of Africa [31,41]. Even though no evidence was highlighted, these changes could be related to alterations in the expression of genes involved in the mosquito visual system.

Another possible implication of the DGK gene in the adaptation of vector populations is its indirect implication to insecticide resistance via the regulation of the acetylcholine release at synaptic junctions. In *C. elegans*, the DGK gene regulates the amount of acetylcholine in synaptic junctions and its inactivation causes a high susceptibility to aldicarb, a carbamate insecticide [43]. This report clearly showed the potential implication of the DGK gene in organophosphates and carbamates resistance *C. elegans*. If synaptic junctions are regulated in the same way in mosquitoes, any selective variant in the DGK gene would affect the acetylcholine release in synapses. These two hypotheses of the potential implication of the DGK gene in adaptations, either insecticide resistance or biting time change, were previously drawn up to explain the selection signal observed in the DGK gene [50]. These hypotheses seem different in both organisms, but the molecular mechanisms underlying the DGK gene function remain similar. Given this intriguing role of DGK, further studies are needed to deeply investigate the role of the DGK gene (AGAP000519) in modulating malaria vector behavior and its potential involvement in behavioral and genetic resistance.

## Conclusion

Investigation of genome variation in vector populations is crucial in understanding dynamics of populations in a given area and guiding strategies of malaria control programs. This study showed a weak differentiation and strong genetic connection between *An. gambiae* and *An. coluzzii* populations in the western area of Burkina Faso. The emergence of new variants together with the already existing pyrethroids resistance variants at high frequencies raise concerns about the long-term efficacy of new-generation nets impregnated with pyrethroids compounds. The long-term efficacy of pirimiphos-methyl-based IRS is also threatened because of the increasing frequency of the SNP *Ace1-G280S* and the duplication of the *Ace1* gene, which confer high fitness to vector populations carrying them. Genome-wide selection scans detected strong signals of positive selection in the genomic regions having an implication in insecticide resistance. This study highlighted the benefit of the molecular surveillance of malaria vectors in the detection of new insecticide resistance variants, the monitoring of the existing resistance variants and also to get insights in evolutionary processes underlying these variants.

## Methods

### Sample collection

Mosquito samples were collected by the Target Malaria project in three sites, Bana (11.233, −4.472), Souroukoudinga (11.235, −4.535) and Pala (11.150, −4.235), located in the western part of Burkina Faso (see Results - Figure 1). These villages are located in the humid savannah zone surrounding the Bobo-Dioulasso city and are characterized by two seasons: a rainy period from May to October and a dry season from November to April. The average annual rainfall in this area is above 800mm (average annual temperature ∼ 27 °C and humidity ∼ 60%) [6]. The main agricultural practices are cotton and cereal (maize, rice, sorghum etc.) cultivation utilizing a large amount of pesticides [53].

Malaria remains endemic in the area surrounding the Bobo-dioulasso city, causing 338 deaths in the three health districts of the city [10]. *An. gambiae* complex species remains the major malaria vector in the sampling area [6]. Malaria control relies on the antimalarial drugs and prevention of mosquito bites using long-lasting insecticidal nets (LLINs), the most widespread vector control tool, and to a lesser extent indoor residual spraying (IRS), sprays and mosquito-repellent coils.

Mosquitoes were collected during the rainy and dry seasons from 2012 to 2017 using three collection methods: Pyrethroid spray catches (PSC), human landing catches (HLC) and swarm collection [54]. PSC and HLC were used for female mosquito collection, whereas males were collected in swarms. After collection, mosquito samples were morphologically identified using the morphological keys [55] and *An. gambiae s.l.* specimens were stored in 80% ethanol and then shipped to Wellcome Sanger Institute for the whole genome sequencing.

### Whole genome sequencing and genomic data management

*An. gambiae* s.l. mosquitoes were sequenced individually at high coverage (30x) using Illumina technology at the Wellcome Sanger Institute. Genomic data were analyzed at the MalariaGEN Resource Centre Team using a wide range of genome analysis software. Reads were aligned to the AgamP4 reference genome using BWA version 0.7.15. Indel realignment was performed using GATK version 3.7-0 RealignerTargetCreator and IndelRealigner. Single nucleotide polymorphisms were called using GATK version 3.7-0 UnifiedGenotyper. Genotypes were called for each sample independently, in genotyping mode, given all possible alleles at all genomic sites where the reference base was not “N”. Coverage was capped at 250× by random down-sampling. Complete specifications of the alignment and genotyping pipelines are available from the malariagen/pipelines GitHub repository [56]. After variant calling, the raw sequences in FASTQ format and the aligned sequences in BAM format were stored in the European Nucleotide Archive (ENA, Study Accession n° PRJEB42254). The genetic variants in VCF and zarr formats, including the samples’ metadata, were stored on Google Cloud and are accessible via the malariagen-data package or are directly downloadable. More details regarding the sequencing technology, the genomic data storage and management including the SNPs variants calling, haplotypes phasing, and copy number variants identification are available on the homepage of MalariaGEN (https://malariagen.github.io/vector-data/ag3/methods.html).

### Data analyses

Genetic diversity and population structure were analyzed in the python programming language on the Google Colab platform. The genetic variants (SNPs, haplotypes and CNVs) and the reference genome of *An. gambiae* s.l. were accessed using the malariagen-data package for direct analyses of the genomic data in the cloud. Specific python packages including malariagen-data [57], scikit-allel [58] and other python standard data-management packages were used for the data analyses. The diversity statistics (segregating sites, nucleotide diversity (*θ_π_*), Tajima’s D (D), Watterson theta (*θ_w_*), allele frequency spectrum) were computed in the whole chromosome in all the mosquito populations. Tukey multiple comparison test was applied to compare the diversity statistic between vector populations over the year. Genome wide selection scan was performed using the Garud h12 [59] statistic to detect the genomic region under positive selection. *F_ST_* and PCA analyses were performed in each chromosome to investigate the genetic structure and the differentiation of *An. gambiae* s.l. populations. The extent and the spread of insecticide resistance variants were also investigated in the *An. gambiae* s.l. populations using the malariagen_data python package. Time series (from 2012 to 2017) SNPs frequencies were analyzed in known target-site resistance associated genes, *Vgsc, Rdl* and *Ace1*. Also copy number variation (CNV) frequencies were investigated in genomic regions associated with metabolic resistance, cytochrome P450s, glutathione-s-transferases, acetylcholinesterase, diacylglycerol kinase and carboxylesterases.

## Ethics approval and consent to participate

All methods in this paper have been implemented in accordance with the relevant guidelines/regulations/legislation in Burkina Faso. No ethics approval was required to run all the activities related to this paper.

## Consent for publication

Not applicable.

## Availability of data and materials

Jupyter Notebooks and scripts to reproduce all the analyses, tables and figures are available in the GitHub repository: https://github.com/mkient/BF_Ag1000G_analyses. The SNPs and haplotypes data are available on the homepage of MalariaGEN and can be accessed using the malariagen-data package.

## Competing interests

The authors declare no competing interests.

## Funding

The sample collection is supported by Target Malaria, which receives core funding from the Bill and Melinda Gates Foundation and Open Philanthropy. Whole genome sequencing and genomics data management were supported by MalariaGEN at the Wellcome Sanger Institute.

## Authors’ contributions

MK, CSC and AM conceived the study. AB and AD provided the resources. AB, AMGB and AD supervised the study. AAM, NT, PSA, SOL and FYA carried out samples collection in the field. AM, CSC and JB produced the genomic data. MK, CSC, HTY and JB carried out data analysis and visualization. MK and CSC drafted the manuscript. All authors have read and approved this version of the manuscript.

## Acknowledgements

The authors would like to acknowledge the Institut de Recherche en Sciences de la Santé, its funders and partners for supporting the sample collection. We are grateful to the Ag1000G project, the MalariaGEN, Sanger Core and GSU Vector Surveillance Programme at the Wellcome Sanger Institute, and their partners in Africa for the production of the *An. gambiae* genomic data. The authors also thank the populations of the sampling sites for their sincere cooperation during the mosquito sampling.

## Supplementary information

**<u>Table S1.</u>** Non-synonymous SNPs frequencies of the *Vgsc* gene (2L: 2358158 - 2431617) within *An. gambiae* s.l. populations.

**<u>Table S2.</u>** Copy number variations of the cytochrome p450 genes and their frequencies within *An. gambiae* s.l. populations.

**<u>Table S3.</u>** Non-synonymous SNPs frequencies of the *Ace1* gene (2R: 3484107 - 3495790) within An. gambiae s.l. populations.

**<u>Table S4.</u>** Copy number variations of the glutathione-s-transferase genes and their frequencies within *An. gambiae* s.l. populations.

**<u>Table S5.</u>** Copy number variations of the carboxylesterase genes and their frequencies within *An. gambiae* s.l. populations.

**Fig. S1.**
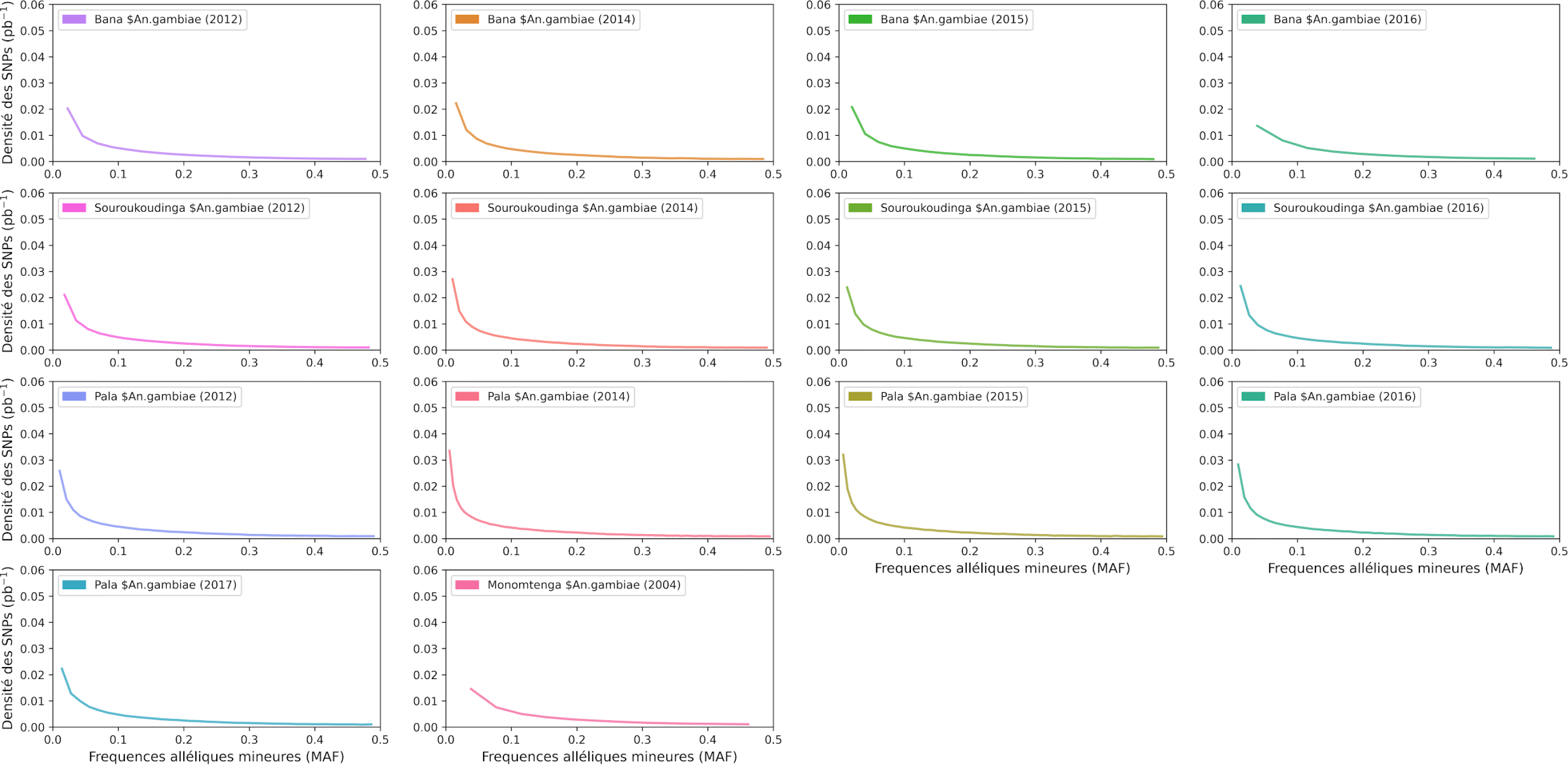
Site frequency spectra in the 3L chromosome of the *An. gambiae* s.s. populations from 2012 to 2016 in the three villages.

**Fig. S2.**
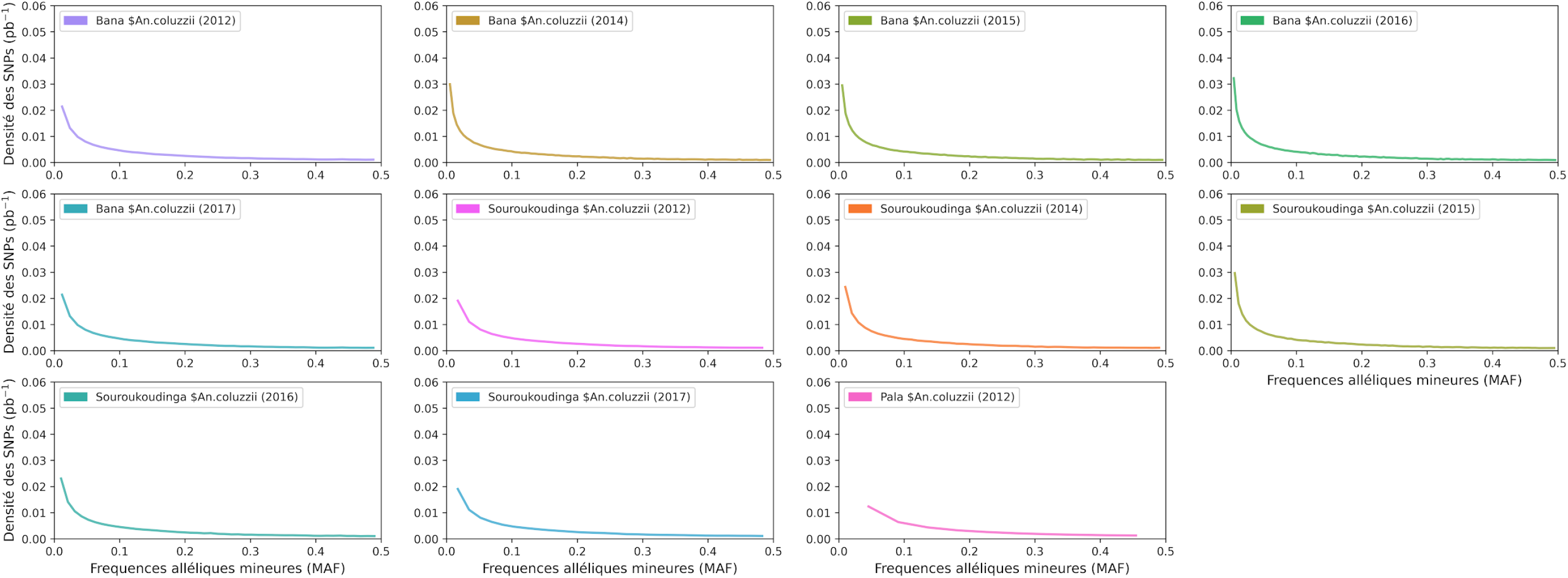
Site frequency spectra in the 3L chromosome of the *An. coluzzii* populations from 2012 to 2017 in the three villages.

**Fig. S3.**
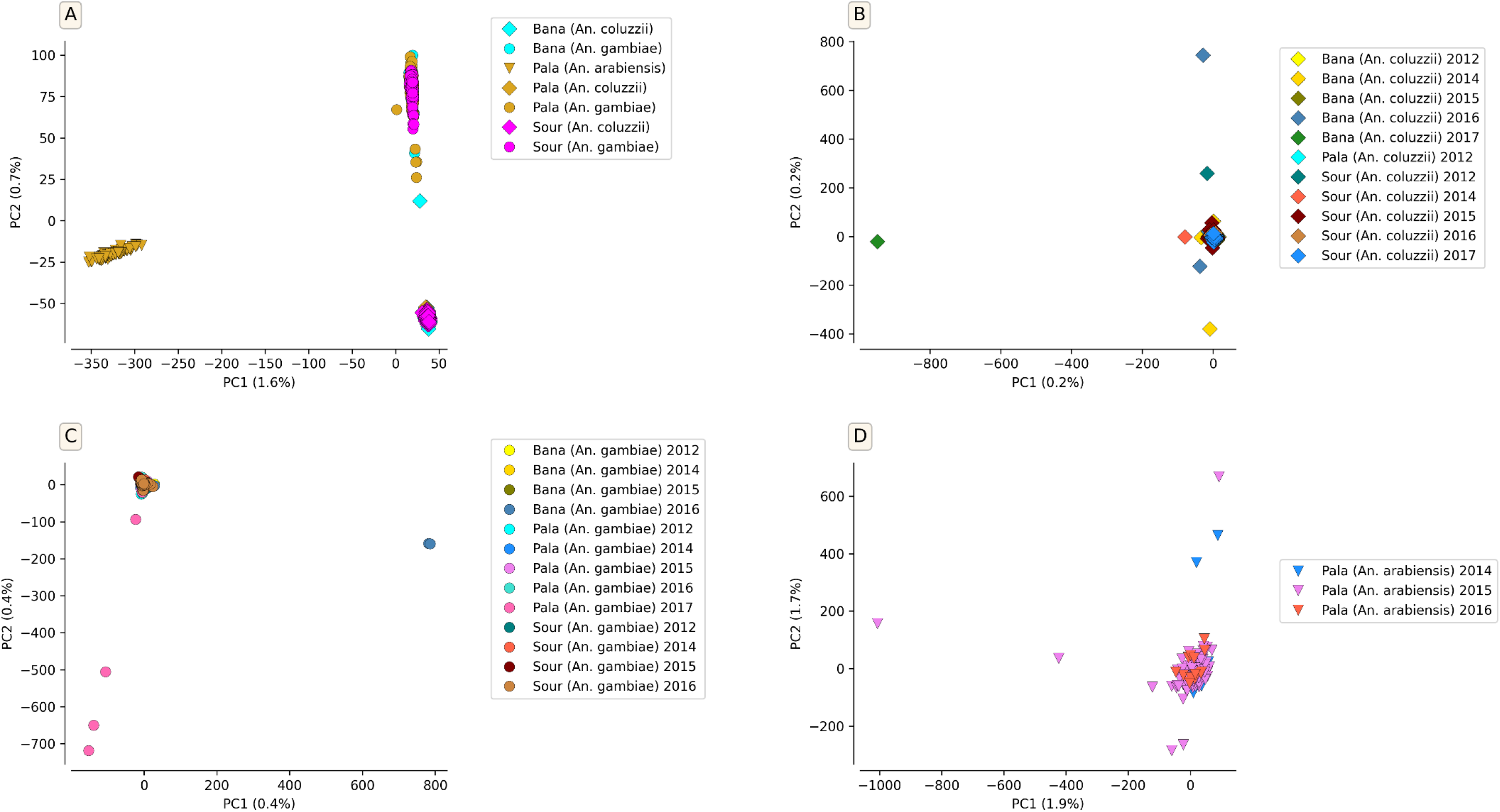
PCA showing the year-to-year (from 2012 to 2017) genetic structure of *An. gambiae* s.l. populations using the whole SNPs identified in the 3L chromosome. B. Year-to-year genetic structure of *An. coluzzii* populations; C. Year-to-year genetic structure of *An. gambiae* s.s. populations; D. Year-to-year genetic structure of *An. arabiensis* populations; These figures demonstrated the lack of geographic substructure within each species of the *An. gambiae* s.l. populations collected in different villages over the years.

**Fig. S4.**
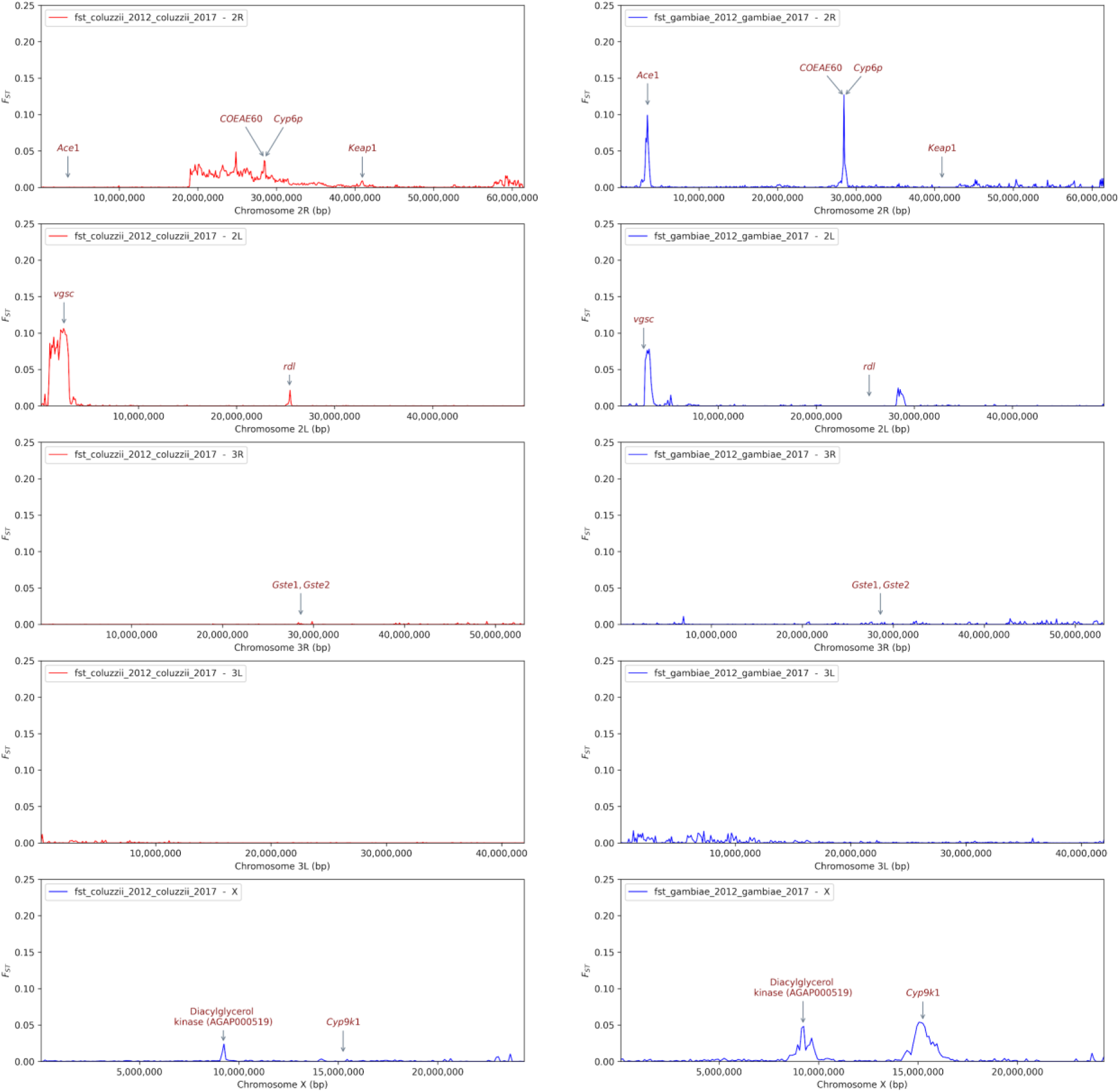
Year to year variation of the genetic differentiation between populations of the same species of *An. gambiae* complex collected in 2012 and in 2017. Signals of differentiation were observed in some regions involved in insecticides resistance. *Ace1*: acetylcholinesterase gene, *Cyp6p*: Cytochrome P450 gene, *Keap1*: Kelch-like ECH-associated protein 1, *Vgsc*: voltage-gated sodium channel, *rdl*: resistance to dieldrin gene, *Gste1*: glutathione S-transferases epsilon 1, *Gste2*: glutathione-S-transferases epsilon 2, *Cyp9k1*: Cytochrome P450 gene.

**Fig. S5.**
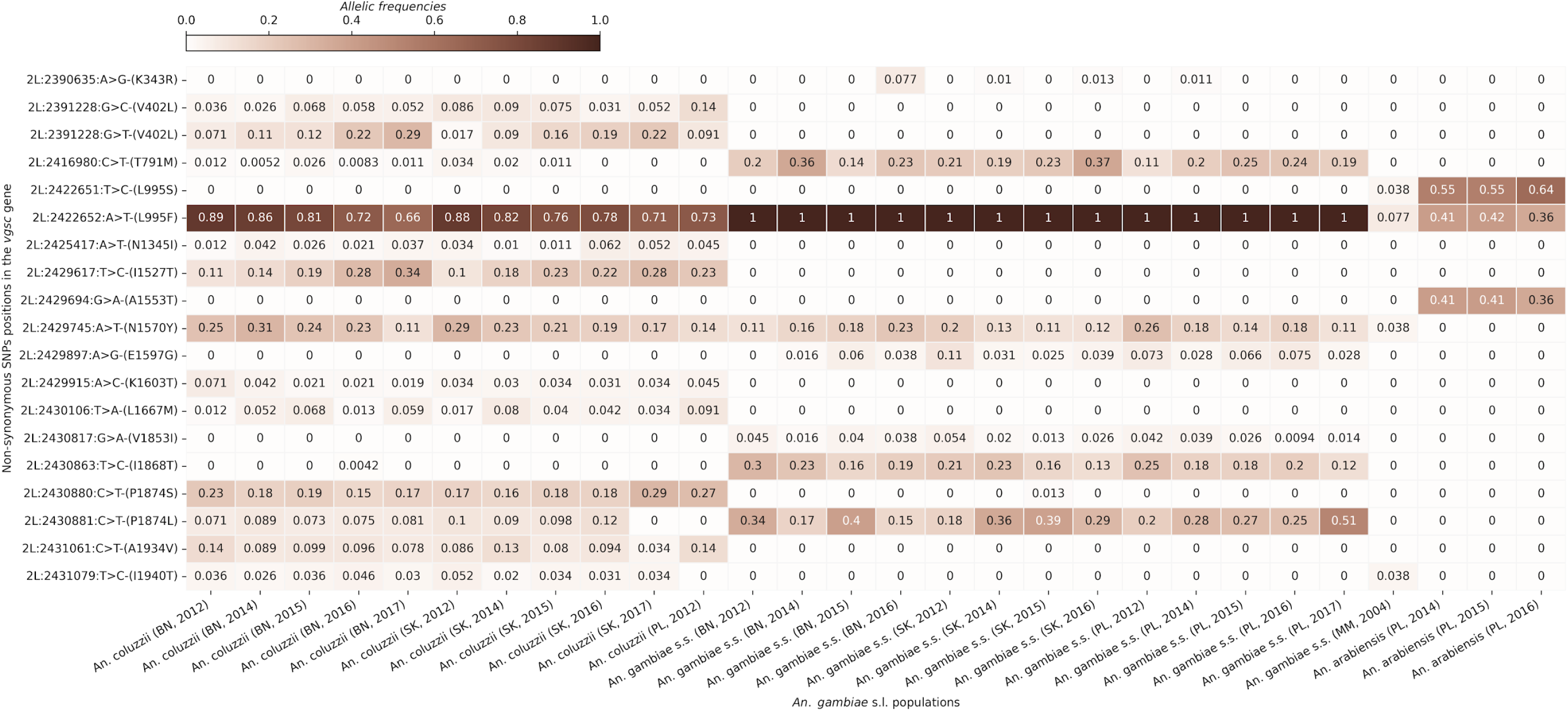
Heat map showing the evolution of the non-synonymous SNPs frequencies (max freq > 0.05) in the *Vgsc* gene within the *An. Gambiae* s.l. populations and the sampling sites. The X axis shows the *An. gambiae* s.l. populations and the years of collection. The Y axis shows the non-synonymous SNPs positions in the chromosome 2L. The gradient color bar shows the distribution of the allelic frequencies. **BN:** Bana, **SK**: Souroukoudinga, **PL**: Pala, **MM**: Monomtenga.

**Fig. S6.**
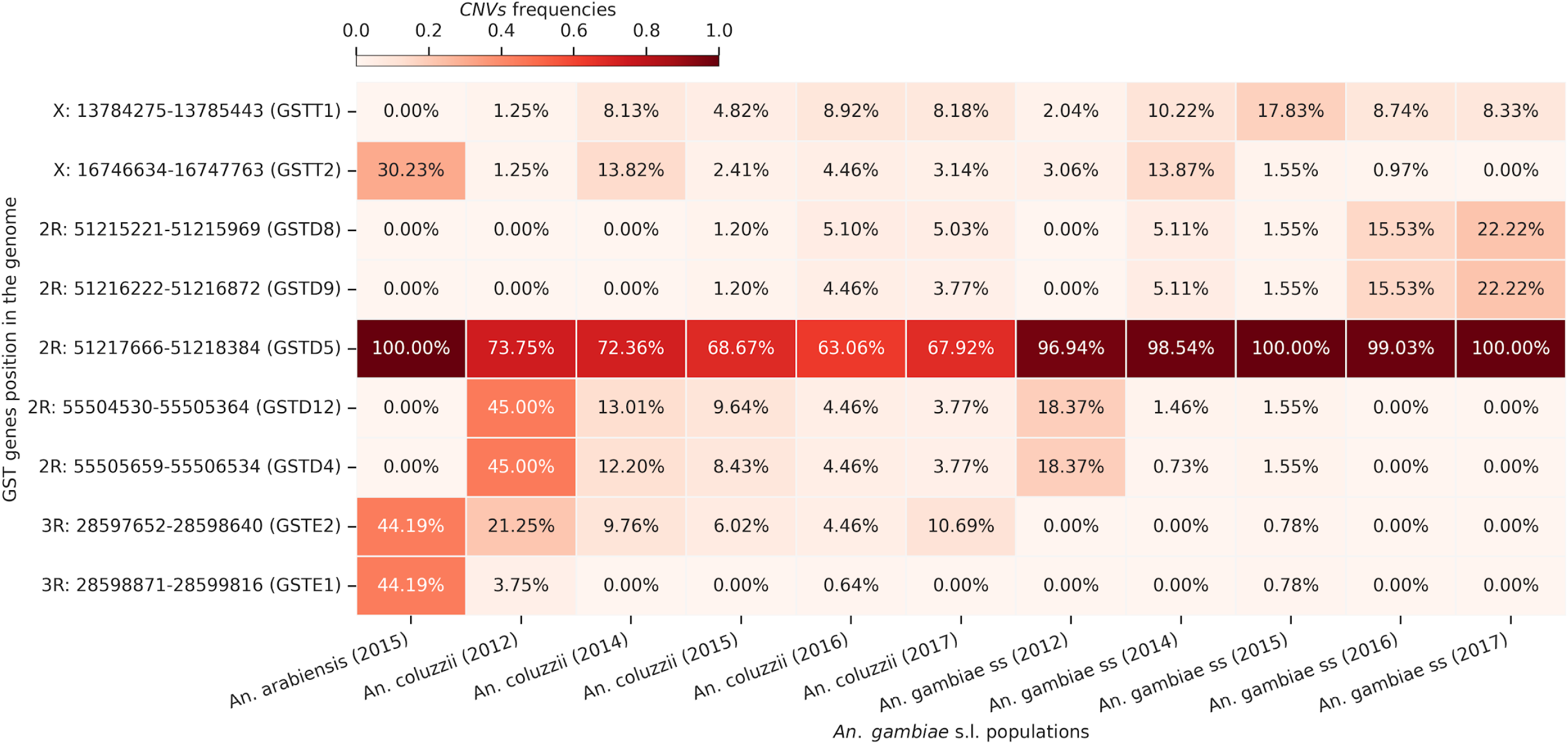
Heat map showing the CNVs frequencies (max freq > 0.05) of the glutathione-s-transferase genes in the *An. gambiae* s.l. populations. The X axis shows the *An. gambiae* s.l. populations and the years of collection. The Y axis shows the positions of the glutathione-s-transferase genes in the genome. The gradient color bar shows the distribution of the allelic frequencies.

**Fig. S7.**
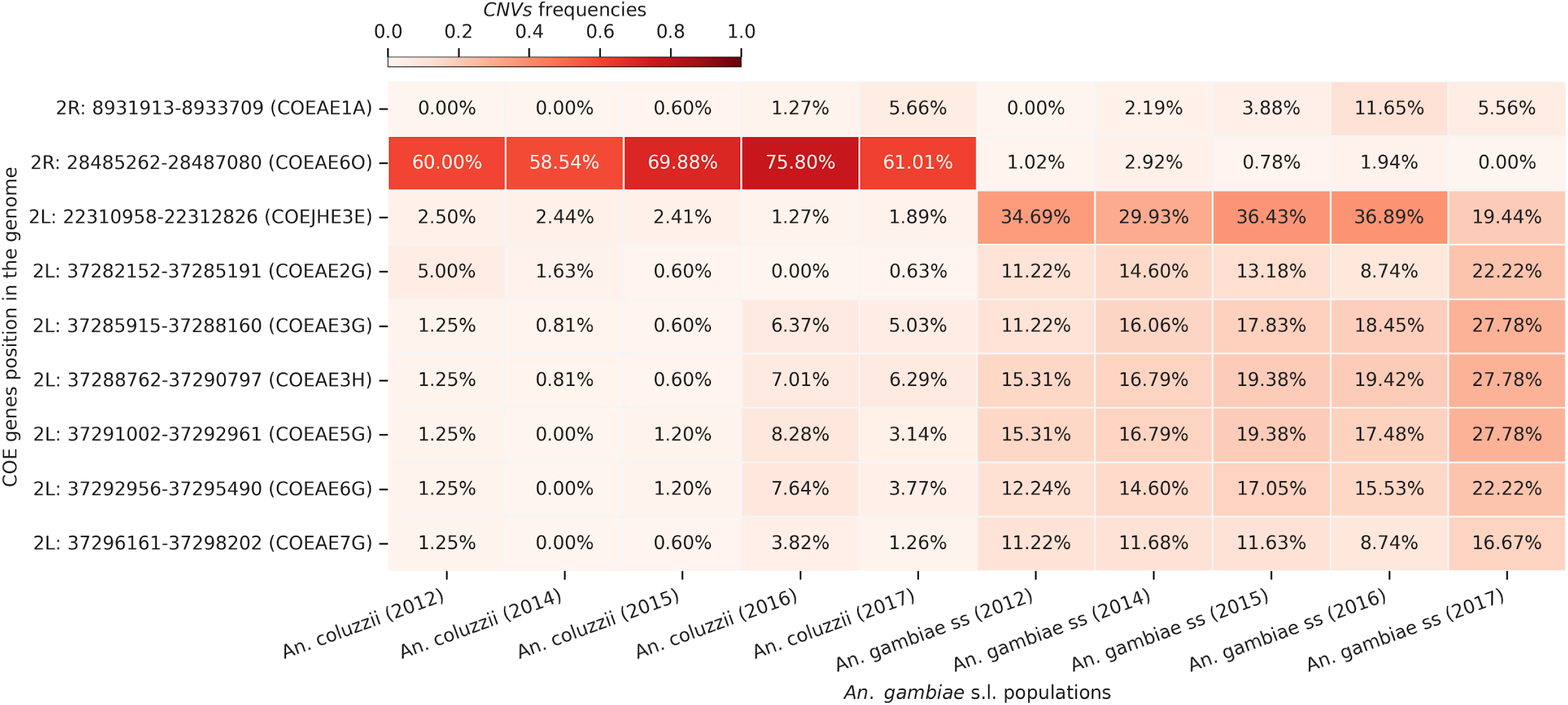
Heat map showing the CNVs frequencies (max freq > 0.05) of the carboxylesterase genes in the *An. gambiae* s.l. populations. The X axis shows the *An. gambiae* s.l. populations and the years of collection. The Y axis shows the positions of the carboxylesterase genes in the genomes. The gradient color bar shows the distribution of the allelic frequencies.

## Notes

### Competing Interest Statement

The authors have declared no competing interest.

### Summary of Updates

Manuscript title revised; Author emails revised; Author contributions revised; References revised;

